# Immunoproteasome deficiency leads to sustained pancreatic injury and delayed recovery from experimental pancreatitis

**DOI:** 10.1101/2020.11.17.386482

**Authors:** Laura L. De Freitas Chama, Frédéric Ebstein, Birthe Wiesrecker, Preshit R. Wagh, Frank U. Weiss, Heike Junker, Maja Studencka-Turski, Markus M. Lerch, Elke Krüger, Matthias Sendler

## Abstract

**Background & Aims:** Uncovering potential new targets involved in pancreas recovery may permit the development of new therapies and improvement of pancreatitis treatment. One disease mechanism comprises the endoplasmic reticulum stress response and a key regulator to prevent proteotoxic stress in an inflammatory context is the immunoproteasome, an induced form of the constitutive proteasome. Our aim was therefore to investigate the role of the immunoproteasome in acute pancreatitis.

**Methods:** Acute pancreatitis was induced in wild type and LMP7^−/−^ mice and several biochemical parameters for disease severity were addressed, including protease activities and histology of pancreatic damage. Real-time PCR was used to measure pro-inflammation and unfolded protein response. Serum IL-6 was detected by cytometric bead assay. Western blotting was used to quantify the ratio of ubiquitin-protein conjugates as well as unfolded protein response activation. Immunofluorescence identified leukocytes infiltration as well as ubiquitin-protein conjugates in the pancreas.

**Results:** In this study, we demonstrate that the β5i/LMP7-subunit deletion correlates with persistent pancreatic damage. Interestingly, immunoproteasome-deficient mice unveil increased activity of pancreatic enzymes as well as higher secretion of Interleukin-6 and transcript expression of the Interleukin IL-1β, IFN-β cytokines and the CXCL-10 chemokine. Thereafter, acinar cell death was increased, which appears to be due to the greater accumulation of ubiquitin-protein conjugates and prolonged unfolded protein response.

**Conclusions:** Our findings suggest that the immunoproteasome plays a protective role in acute pancreatitis via its role in the clearance of damaged proteins and the balance of ER-stress responses in pancreatic acini as well as in macrophages cytokine production.

## Introduction

Acute pancreatitis, one of the most common gastrointestinal diseases, is an inflammation of the pancreas, thought to be primarily due to premature intra-acinar activation of digestive pancreatic zymogens (1–2). Although most patients present with a mild form of the disease, about 20% develop severe pancreatitis associated with organ dysfunction, requiring intensive care (3–4). The global incidence of acute pancreatitis has been reported by Xiao et. al. (5) to be 34 cases per 100.000 per year. Autodigestion of the pancreas by its own proteases leads to acinar cell injury and subsequently to a local and systemic inflammatory response (6–7). Nevertheless, paradoxical results from trypsinogen deficient mice, which still developed experimental pancreatitis (8–9), has brought this traditional concept under review and suggest that the molecular mechanisms associated with the onset disease and progression are not fully understood. Many studies, specifically in animal models, have identified a variety of cellular events as being involved in the pathogenesis, such as premature activation of pancreatic enzymes (10), increase in calcium signaling (11–12), inflammatory cell infiltration (13–14), mitochondrial dysfunction (15) and endoplasmic reticulum stress (ER stress) (16–18).

Acini are the major source of digestive enzymes, thereby exhibiting the highest rate of protein synthesis and folding capacity with more abundant ER then all other cell types (19). ER stress is caused by accumulation of misfolded or unfolded proteins, which may arise during synthesis, folding and secretion of secretory and cell-surface proteins. Therefore, elucidating the molecular mechanisms of protein homeostasis may identify potential new targets of acinar cell injury. ER stress is counterbalanced by the unfolded protein response (UPR) machinery, which has a dedicated role in protein quality control and homeostasis (20). Impairment of UPR was found to be one cause of genetic varieties of pancreatic disorders (21–22).

The ubiquitin proteasome system (UPS) plays an important role in intracellular protein degradation and turnover in eukaryotes via a multi-enzymatic machinery, entailing target protein ubiquitination and subsequent proteolysis by the 26S proteasome (23). The immunoproteasome is an isoform of the constitutive proteasome (26S proteasome), which is assembled upon a triggering inflammatory response by Toll-like or cytokine receptor signaling, type I or type II interferons (IFNs), and TNF-α releases. It arises from the replacement of the β-catalytic subunits β1, β2 and β5 in the 20S standard proteasome core by iβ1 (large multifunctional peptidase 2, LMP2 encoded by PSMB9), iβ2 (multicatalytic endopeptidase complex-like-1, MECL-1 encoded by PSMB10) and iβ5 (LMP7 encoded by PSMB8), respectively, during de novo proteasome formation (24–25). Over the years the immunoproteasome has been recognized as an important player in shaping innate and adaptive immune responses by degradation of inflammatory mediators and improved MHC class I antigen presentation (26–27).

In addition to its roles in innate and adaptive immune cells, the immunoproteasome has also been linked to non-immune functions, namely cell differentiation and protein homeostasis, including intracellular protein clearance (28) and subsequently, may thus control of the ER stress levels. Likewise, a growing body of evidence has unveiled a consistent interdependence between ER stress imbalance and the immune system (29–30). In view of the immunoproteasome role for protein clearance and ER-stress pathways, and the association of human mutations with autoinflammatory diseases (31), our aim was to investigate the role of the β5i/LMP7 subunit in pancreatitis. We found that the immunoproteasome has a protective role in acute pancreatitis, since β5i/LMP7 deletion increased disease severity and impeded recovery. Our work demonstrates that this proteasome isoform is involved in the clearance of ubiquitin-protein conjugates, attenuation of the pro-inflammatory response and UPR in pancreatitis.

## Results

### The β5i/LMP7 subunit is upregulated in acute pancreatitis

First, we determined whether β5i/LMP7 subunit and its constitutive counterpart β5 were expressed and differentially modulated in an *in vitro* model of pancreatitis using isolated acini. After supramaximal stimulation of the cells for 8h and 24h with the gastrointestinal hormone CCK, we assessed the protein levels of both subunits by western blotting. Pancreatic acini express basal levels of both subunits but only the β5i/LMP7 subunit was upregulated after 24h of treatment (Fig. 1A). In a next step we investigated whether the regulation of the β5i/LMP7 subunit would also be shifted in an *in vivo* mouse model of pancreatitis. We therefore used the caerulein model, which induces a mild and reversible form of the disease (33). Three experimental groups were designed: control (0h), acute phase (8h) and recovery phase (24h). Interestingly, the β5i/LMP7 transcript levels were significantly increased at 8h after the onset of pancreatitis. At 24h its levels returned to baseline as measured in the control group (Fig. 1B). Importantly, no β5i/LMP7 transcripts were detected in the pancreas from β5i/LMP7^−/−^ mice. Western blot analysis of pancreas homogenates showed an approximately threefold increase of β5i/LMP7 protein levels at 24h compared to basal levels (Fig. 1C).

**Figure 1:**
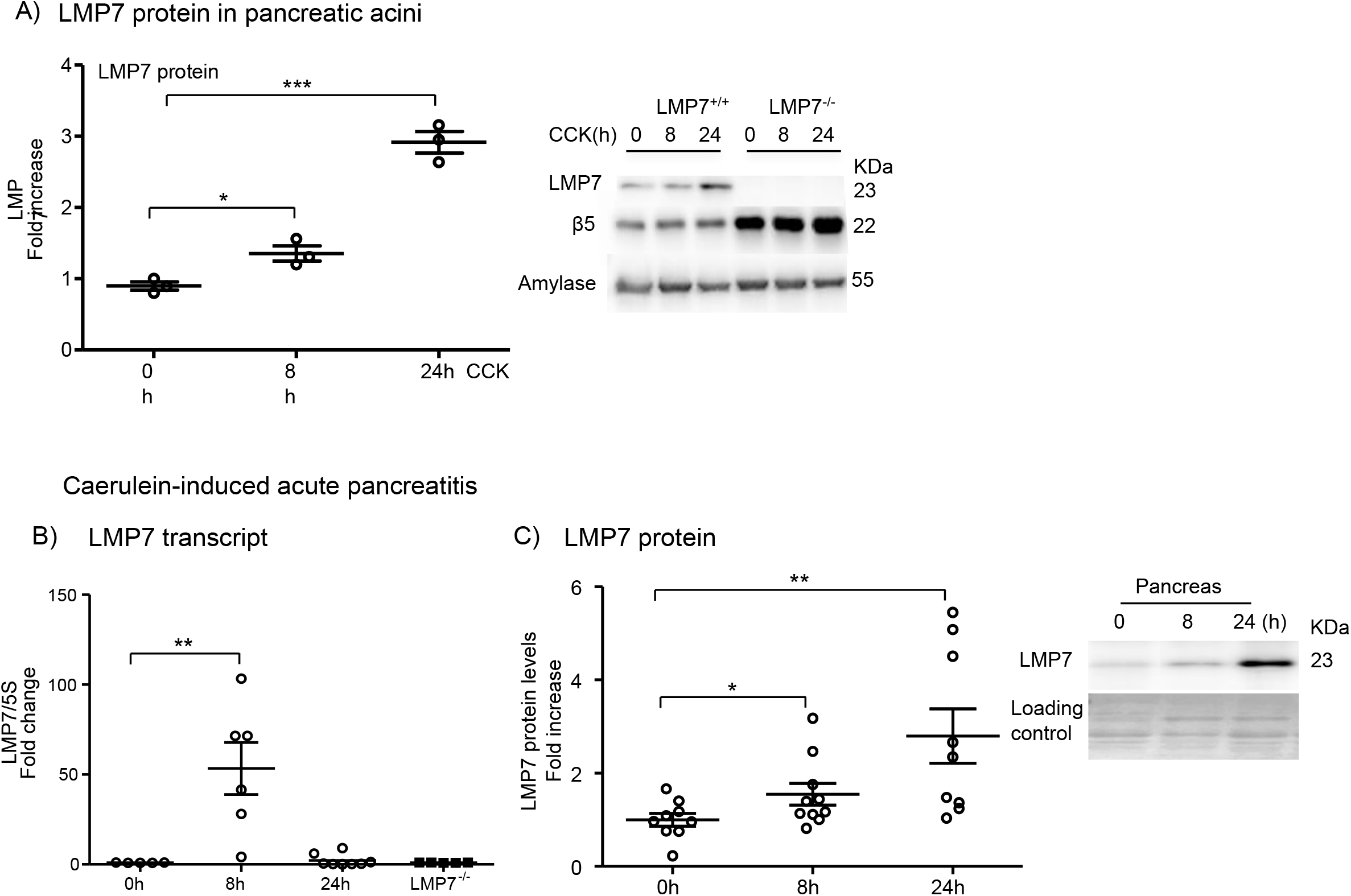
Expression of LMP7 subunit in acute pancreatitis.

### Higher pancreatic damage in β5i/LMP7-deficient mice

To investigate the role of the β5i/LMP7 immunoproteasome subunit in pancreatitis, suggested by its up-regulation *in vivo*, disease severity was evaluated in β5i/LMP7^−/−^ and wild-type littermate β5i/LMP7^+/+^ mice. We performed the caerulein-induced pancreatitis model, including the same experimental groups as before. Biochemical markers of pancreatic injury (13), such as serum amylase and lipase activities, were increased by about 20% in β5i/LMP7^−/−^ mice (Fig. 2A). Additionally, we measured trypsin and chymotrypsin activities in the pancreas homogenates of littermate and β5i/LMP7^−/−^ mice. Trypsin and chymotrypsin consistently showed higher activities in the absence of β5i/LMP7 (Fig.2B). To examine early protease activation of living acini towards stimulation in the absence of β5i/LMP7, we performed an *ex-vivo* enzymatic assay (34). In the time-course of 1h CCK treatment, activities of trypsin and cathepsin B were similar in both genotypes, as well as the necrotic rate (Suppl. Fig.1). Thus, we showed that the lack of β5i/LMP7 did not alter the early protease activation within acini but during later pancreatitis progression.

**Figure 2:**
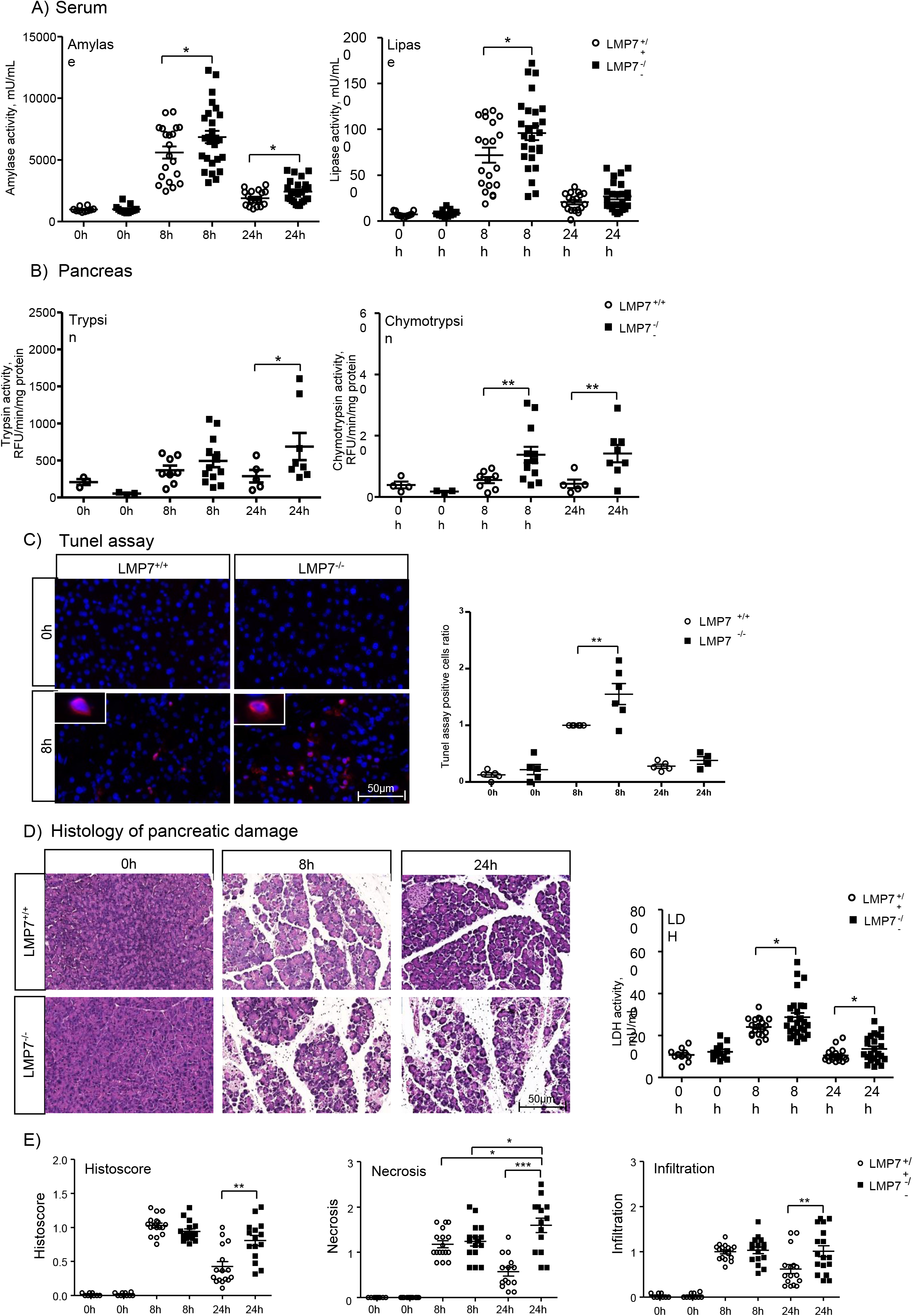
Higher pancreatic damage in the absence of LMP7.

Since intracellular protease activation is associated with acinar cell death (34), we next investigated whether apoptosis and necrosis were likewise increased in β5i/LMP7^−/−^ mice. Apoptosis quantification was accomplished by labeling DNA fragmentation (Tunel assay) in paraffin-embedded pancreas. We observed that β5i/LMP7^−/−^ mice exhibited a significantly higher number of Tunel-positive staining, approximately 1.5 fold higher at 8h, compared to littermate controls (Fig.2C). Histological pancreatic damage was further verified by hematoxylin and eosin staining (13), allowing us to identify and score necrotic areas and leukocytes infiltration. Pancreas from wild-type and β5i/LMP7^−/−^ mice exhibited equivalent pancreatic damage in the acute phase of the disease (8h). However, necrosis was increased at 24h in β5i/LMP7-deficient mice and the tissue damage persisted for a longer period compared to the littermate group (Fig. 2D and E). To assess cell death, we performed serum lactate dehydrogenase activity assay, which was also increased by 20% in β5i/LMP7^−/−^ mice at 8h (Fig.2D). Pancreatic injury was approximately two-fold higher in the absence of β5i/LMP7 at the 24h time point (Fig. 2E), implying a role of the immunoproteasome in pancreas recovery after pancreatitis. Of note, increased inflammatory cell infiltration was detected in the absence of β5i/LMP7 (Fig. 2E), leading the next question to be addressed.

### β5i/LMP7 deficiency correlates with increased inflammation

Our group has previously shown that acinar cell damage is linked to a pro-inflammatory state with increased infiltration of macrophages and neutrophils into the pancreas in response to cell injury (38). We therefore next characterized these leukocytes population by immunofluorescence in pancreatic tissue from all experimental groups. This approach was performed using specific antibodies for profiling of pro- and anti-inflammatory macrophages phenotypes (CD68 and CD206, respectively) and neutrophils (Ly6g). No significant difference was noticed in the count of infiltrating macrophages (Fig. 3A). However, a significant increase in neutrophil population was observed at 24h (Fig. 3B). Next, we measured the myeloperoxidase (MPO) activity in lungs, which is a peroxidase enzyme abundantly expressed in neutrophil granulocytes (39). As expected from the neutrophil infiltration in the absence of β5i/LMP7, we also detected enhanced MPO activity (Fig.3B). We then quantified the activation of pro-inflammatory mediators in serum and in pancreas. Along with extended acinar cell injury we found that the transcription of pro-inflammatory cytokines IL-1β, CXCL-10 at 8h and IFN-β at 24h was increased in the absence of β5i/LMP7 compared to littermate controls by approximately 2.5 to 6-fold (Fig. 3C). Likewise, serum levels of circulating IL-6 cytokine were elevated after 8h of pancreatitis (Fig. 3D). To analyze whether the alterations in the pro-inflammatory cytokines regulation originated from pancreatic acini or macrophage BMDM cells, acinar cells were either stimulated with CCK alone or co-incubated with BMDM cells. This second approach allows the evaluation of macrophages activation in the onset of pancreatitis (38). In both cell types, we observed a higher regulation of IL-6 and IFN-β cytokines in the absence of β5i/LMP7. The CXCL-10 chemokine was upregulated in pancreatic acini whereas IL-1β was significantly elevated in BMDM (Suppl. Fig.2). Taken together, our cytokine data indicate an increase in pro-inflammatory cytokines in acute pancreatitis in the absence of β5i/LMP7 subunit.

**Figure 3:**
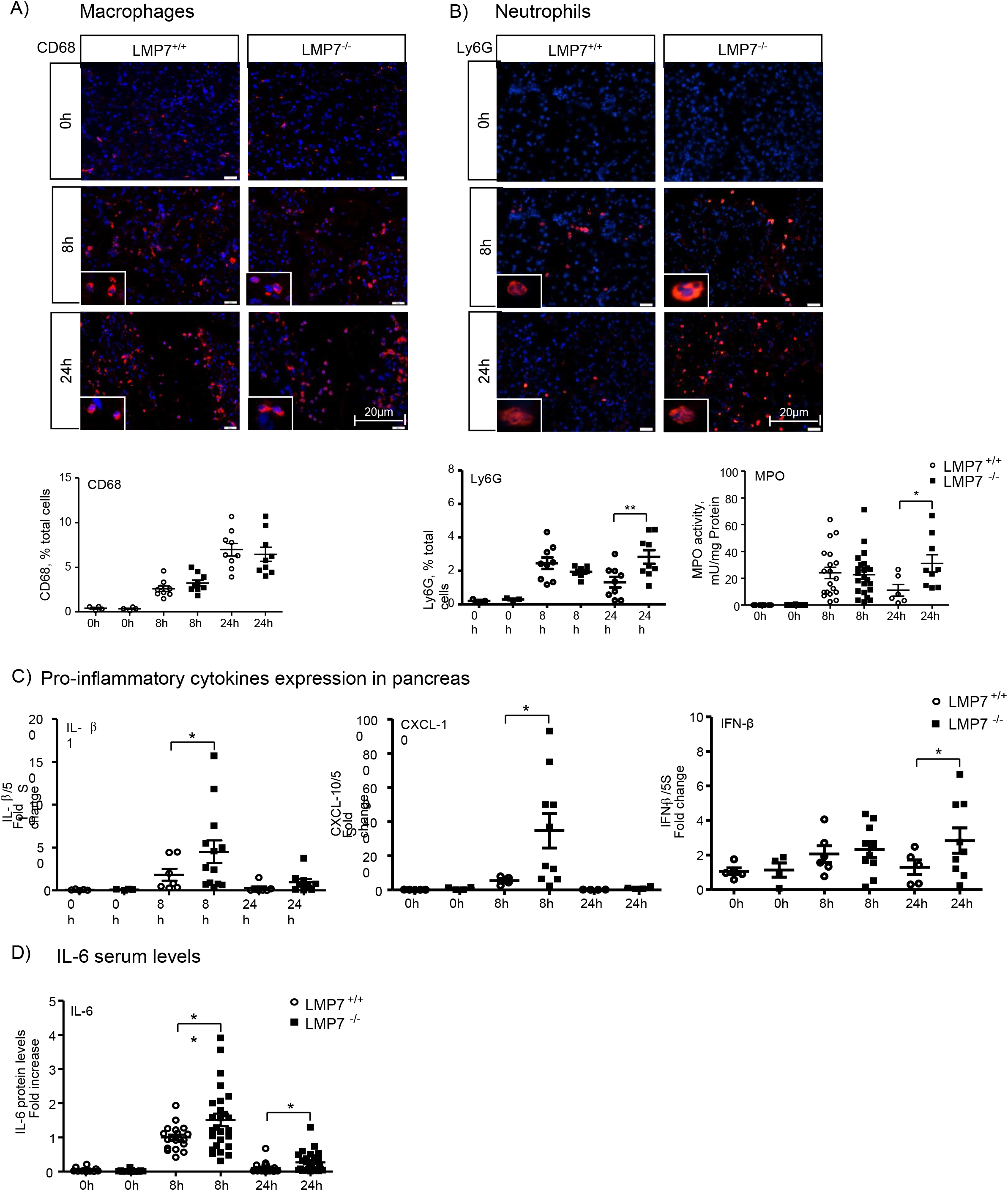
Increased inflammation in the absence of LMP7.

### Impairment of ubiquitinated protein degradation in absence of β5i/LMP7

In order to elucidate the molecular mechanisms associated with the disease phenotype of β5i/LMP7^−/−^ mice during pancreatitis, we examined the immunoproteasome function by immunofluorescence and western blotting; specifically, we analyzed the profile of ubiquitin-protein conjugates. First, we visualized the protein conjugates in the pancreas by immunofluorescence using an anti-ubiquitin antibody. Our data show an increased accumulation of ubiquitin-modified proteins at 8h in the absence of β5i/LMP7, which was quantified by western blotting in insoluble fractions of pancreas homogenates from all experimental groups (Fig. 4A and B). These findings suggest that an impairment of immunoproteasome function has an impact on protein degradation and subsequent proteotoxic stress during pancreatitis.

**Figure 4:**
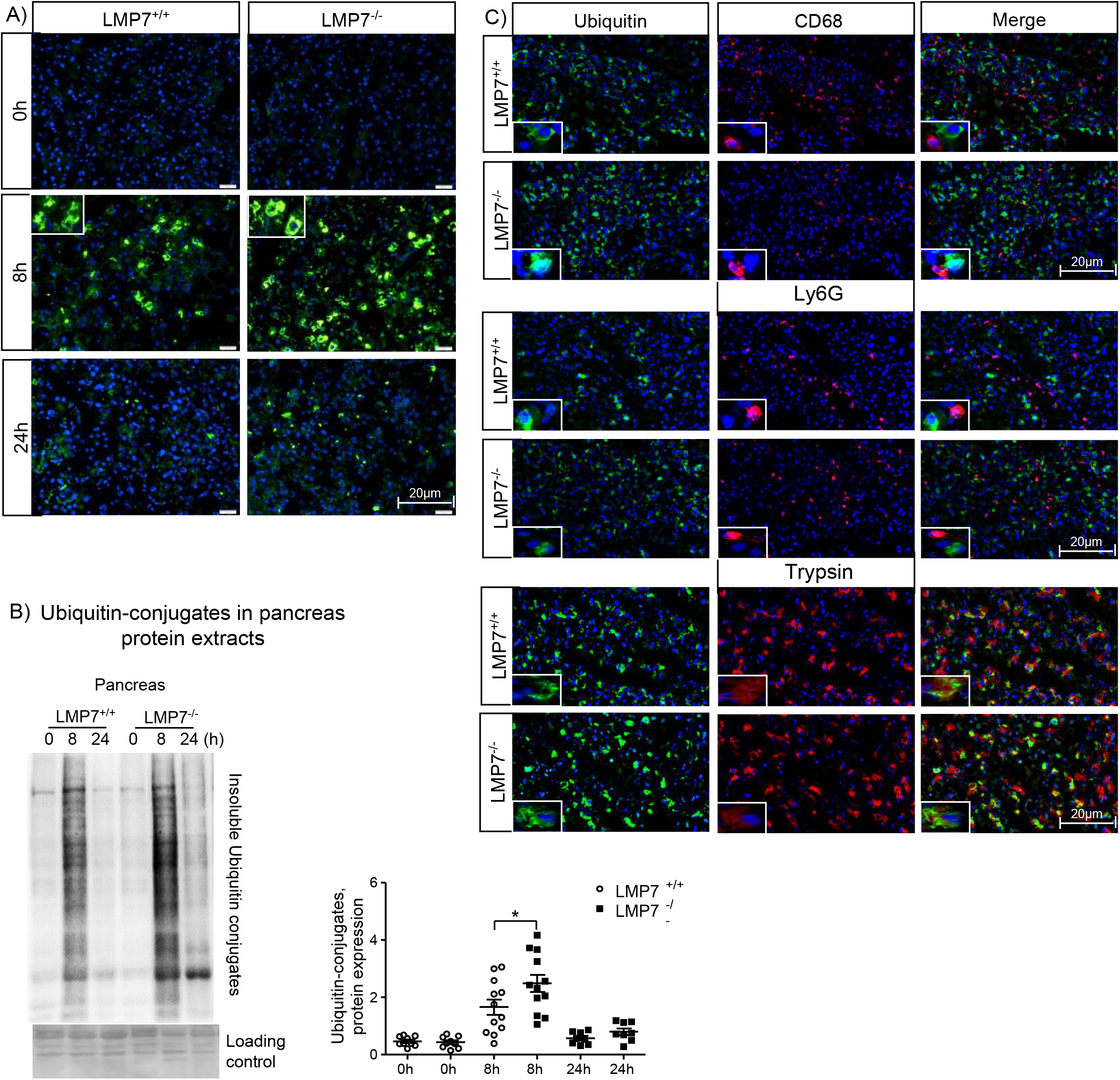
Higher accumulation of ubiquitinated-protein conjugates in pancreatic acini from LMP7-deficient mice.

Our next step was to determine whether the ubiquitin-protein conjugates were localized in pancreatic and/or immune cells by co-immunofluorescence experiments on pancreatic tissue. Strikingly, the conjugates were detected exclusively in acini but not in macrophages or neutrophils (Fig. 4C). In addition to the ubiquitin-dependent protein degradation pathway, we assessed whether the absence of β5i/LMP7 also had an influence on autophagy. Autophagy dysregulation has been identified as one key mechanism involved in the pathogenesis of pancreatitis. LC3-II is the lipidated form of its cytosolic counterpart (LC3-I) and required for autophagosome formation. Sequentially, it engulfs cellular material for degradation and fuses with lysosomes, comprising the entity, in which protein cleavage occurs (40). Analysis of LC3-II protein levels showed similar activation in the absence of β5i/LMP7, inferring that the induction of autophagy was not changed in our model (Suppl. Fig.3) and does not depend on the immunoproteasome.

### Sustained modulation of the unfolded protein response in β5i/LMP7^−/−^ mice

The expected consequences of immunoproteasome impairment would be a delayed protein clearance (41). Thus we hypothesized that immunoproteasome-deficiency may result in increased endoplasmic reticulum stress and consequently, turn-on of the UPR signaling pathways. To address the UPR response, we performed quantitative real-time PCR in the pancreas to determine the expression of BIP and the transcription factors ATF4 and sXBP1, which are sensor and downstream targets of the UPR, respectively (Fig. 5A). LPM7 deficiency had no effect on the transcript regulation in the acute phase (8h) of the disease. Nevertheless, there was a prolonged induction of the transcripts at 24h in the absence of β5i/LMP7, with an up to twofold increase over littermate controls. In addition, protein levels of C/EBP homologous protein (CHOP) (Fig.5B), which is a downstream target of the ATF4 and sXBP1 pathways, were significantly enhanced at 24h in the pancreas of β5i/LMP7^−/−^ mice, confirming the importance of a functional immunoproteasome in regulating UPR induction. Subsequently, the ER stress response was also investigated in *in vitro* experiments using pancreatic acini and co-culture of acini and BMDM (Fig. 5C and D). There was a clear induction of ER stress response transcripts BIP, ATF-4 and sXBP-1 in both cell types, but only acini underwent an altered regulation in the absence of β5i/LMP7.

**Figure 5:**
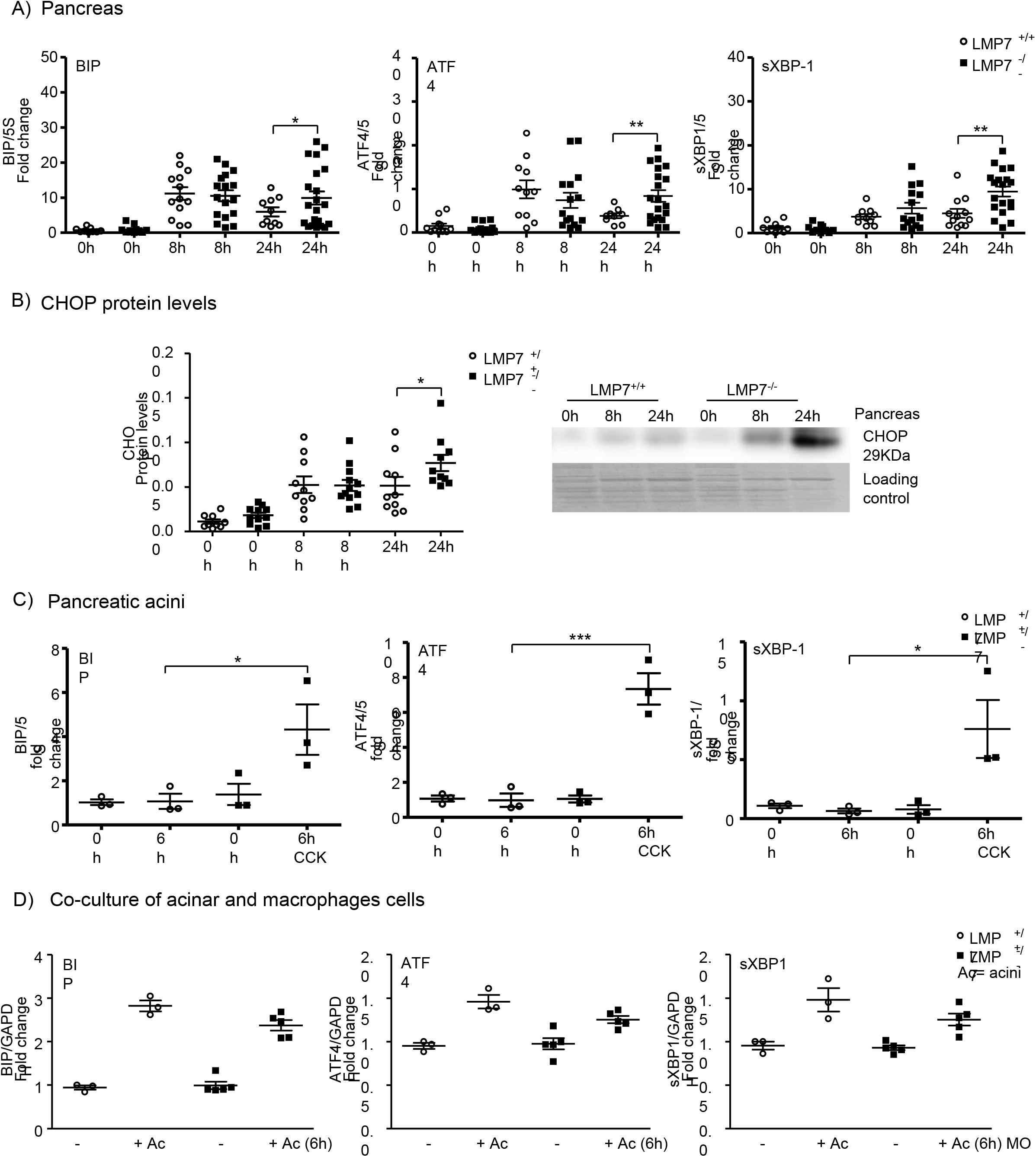
Stronger induction of UPR in the absence of LMP7.

Overall, our data imply that the immunoproteasome plays a protective role in acinar cell homeostasis, since elevated cell death was observed in the absence of β5i/LMP7 in pancreatitis. The effect on pancreas recovery did not relate to changes in acinar cell regeneration, considering that the proliferation rate visualized by immunohistochemistry was not altered in the absence of β5i/LMP7 (Suppl. Fig.4). Therefore, the immunoproteasome appears to contribute to the clearance of ubiquitin-protein aggregates, resolution of inflammation and the reversal of UPR activation in pancreatitis.

## Discussion

In this study we report a protective role of the immunoproteasome in pancreatitis and a beneficial effect on pancreatic injury resolution. The expression of the immunoproteasome subunits is known to be regulated not only in immune cells but also in non-immune cells, for example in pancreatic beta cells, muscle cells and adipocytes (42–44). Previous reports have demonstrated the importance of this proteasome isoform in cell differentiation and protein homeostasis (41,45). Our work provides evidence for a role of the immunoproteasome in pancreatic acinar cell homeostasis and organ repair after cell injury. Notably, the β5i/LMP7 subunit but not the β5 constitutive subunit was up-regulated in an *in vitro* and an *in vivo* model of pancreatitis. Increased β5 protein levels in acini from β5i/LMP7-deficient mice were observed as a compensatory mechanism, as seen before in other cell types (46–47). However, from the data obtained in this study, the immunoproteasome represents the major proteasome type involved in a rapid and efficient response to inflammation and ER stress during pancreatitis.

In the development of pancreatitis, well-known as a primarily sterile disease, two major players have been described in the past. Injured acinar cells release cellular components, for instance free ATP, histones or DNA, which act as damage-associated molecular patterns. These elicit the recruitment of macrophages and neutrophils into the pancreas, which then trigger inflammation and are associated with tissue injury (38). Beyond immune cells, acinar cells *per se* can undergo NF-kB transcription factor activation and secrete pro-inflammatory cytokines and chemokines, such as IL-6, TNF-α and MCP-1 (48). On the other hand, several reports have associated β5i/LMP7 inhibition or deficiency with altered cytokine production (49–52). Here we show an opposite effect in which immunoproteasome deficiency correlates with an increased production of pro-inflammatory cytokines. In this context acini and macrophages were affected, considering that IL-1β is exclusively expressed by macrophages (14). Besides the differences detected in cytokine production, we also observed that immunoproteasome deficiency resulted in higher infiltration of neutrophils into the pancreas. Over the last decade, various studies have found an association between oxidative stress and acinar cell damage, with neutrophils as the central player in this process (53). The difference in neutrophil infiltration may possibly account for the stronger oxidative stress in the absence of β5i/LMP7. In line with this, Seifert et al. have previously shown a primordial role of the β5i/LMP7 subunit in preserving cell viability upon cytokine-induced oxidative stress (41).

The second aspect of our study highlights the effect of the immunoproteasome on acinar cell protein homeostasis. It is well established that acini, in response to supraphysiological secretagogue stimulation, develop ER stress and subsequently activate UPR (54). In turn, UPR upregulation has been recognized as a protective mechanism against acinar cell injury in pancreatitis (16). The UPR restores ER protein homeostasis through inhibition of protein synthesis, enhancement of protein folding and degradation mechanisms, such as autophagy and the UPS (20). Recently, the UPR and the immunoproteasome have appeared as pivotal networks in the maintenance of protein homeostasis and cell fate control (55–56). In our work, we show that the immunoproteasome is crucial for limiting UPR activation in isolated pancreatic acini. The UPR encircles stress response signaling pathways triggered by accumulation of misfolded or damaged proteins in the ER lumen (57). Although some studies did not report differences in the capacities of the constitutive- and the immunoproteasome to degrade ubiquitinated protein targets (47,58), our work supports an essential role of the immunoproteasome in regulating this process and synchronizing UPR activation in pancreatitis. Unbalanced UPR, due to proteasome dysfunction has been implicated in the innate immune response, with the immunoproteasome playing a central role in the regulation of inflammation and UPR. The crosstalk between UPR and type I IFN signature has been proposed to evoke a sterile inflammation in the event of increased ER stress levels (59–62). Here, we also found β5i/LMP7 deficiency to be associated with an altered ER stress response and enhanced IFN and IFN-stimulated genes, such as IFN-β and CXCL-10. We have previously shown that the IFN type I signaling pathway could be a promising target in the pancreatitis immune defense, using macrophages exposed to acini (14). Thus, our work reveals a functional link between IFN type I signaling and the UPR activation in pancreatitis, with the main effect on acinar cell protein homeostasis.

As a consequence of UPR dysregulation, represented by a prolonged time span for activation of sensor proteins and downstream targets, cell death appears to be the outcome (56). As verified by pancreas histology, immunoproteasome dysfunction correlated with sustained pancreatic damage, which was accompanied by increased acinar cell death, namely apoptosis and necrosis. These observations support previous findings, implying that the UPR is to a certain extent protective via restoring protein homeostasis. Nevertheless, its sustained activation may lead to cell death (20,63). One of the proposed mechanisms involves the ATF4-CHOP signaling pathway, downstream targets of the PERK stress sensor. Our data align with previous information regarding the activation of this UPR pathway in acute pancreatitis (16). Beyond this we identified a role of immunoproteasome in preventing deleterious ER stress levels and maintaining acinar cell viability.

Lastly, Zhu and coworkers have demonstrated beneficial effects of a proteasome inhibitor, named bortezomib, in two models of pancreatitis (64). In their work, significant differences in the histological score have been observed with the highest dose of bortezomib, which mitigated the disease severity through inhibition of acinar cell necrosis and suppression of NF-kB activity. The discrepancies to our β5i/LMP7 data are most likely due to the inhibition of all proteasome types by bortezomib.

In conclusion, the balance of ER stress levels and the maintenance of acinar protein homeostasis are important cellular processes in the resolution of tissue injury. In this context, the immunoproteasome appears to be a critical proteolytic complex in mitigating the pro-inflammatory response, preventing pancreatic injury and promoting subsequent recovery from pancreatitis. Future studies will be needed to identify pharmaceutically usable agents that improve or maintain the function of the immunoproteasome in order to translate this novel treatment target into a therapeutic modality.

## Material and Methods

### Animals

Experiments were performed using male and female β5i/LMP7^+/+^ (wild-type littermate) and β5i/LMP7^−/−^ mice C57BL/6J strain background, generated as previously reported (32), weighing 20 to 25g at about 2-3 months of age. Both genotypes were bred and maintained at the Greifswald University Animal Care Facility. Food and water were provided *ad libitum* to the animals, which were kept in a controlled environment with a constant 12:12-h light– dark cycle.

### Caerulein-induced acute pancreatitis model

Acute pancreatitis was induced by eight hourly intraperitoneal injections of caerulein (50μg/kg bodyweight; cat: C9026-Sigma, Merck, Darmstadt, Germany), as previously described (33). The experimental protocol was approved by the institutional Animal Care and Use Committee. Three experimental groups were designed: control animals (0h; no caerulein injections), 8h corresponding to the acute phase of the disease and 24h to the recovery phase. Five independent experiments were performed and no difference regarding sex was statistically significant. Serum and tissues were collected, processed and adequately stored to the experimental procedures, as described ahead.

### Biochemical assays

Quantification of serum activities of amylase and lipase enzymes were assayed by photometric assays kits from Roche Hitachi (Grenzach-Wyhlen, Germany). Lactate dehydrogenase was measured in serum using a specific kit assay from Sigma (cat: MAK066, Merck). Pancreatic enzymatic activities, such as trypsin and chymotrypsin were determined in pancreatic tissue homogenates using fluorescent substrates as previously reported (34): Trypsin substrate (R110-(CBZ-Ile-Pro-Arg)2 was purchased from Life Technologies (Carlsbad, CA, USA); Chymotrypsin (Suc-AAPF-AMC) and Cathepsin B substrates (AMC-Arg2) were obtained from Bachem AG (Bubendorf, Switzerland). Myeloperoxidase (MPO) activity measurement in lung was performed as previously described (13). All activities from tissue homogenates were normalized suitably by protein concentration using Pierce BCA protein assay (cat: 23227, Thermo Fisher Scientific, Waltham, MA, USA).

### Measurement of pro-inflammatory cytokines

The IL-6 pro-inflammatory cytokine was measured in serum and supernatant samples by fluorescence activated cell sorter (FACS) analysis using the CBA mouse inflammation kit, according the manufacturer’s instructions (cat: 552364, Becton Dickinson, Heidelberg, Germany). IL-1β cytokine was measured in supernatant from macrophage culture using Peprotech kit assay (cat: 900-M47, Hamburg, Deutschland).

### Imaging analyses

Histological pancreatic damage was verified by hematoxylin and eosin (H&E) staining in paraffin-embedded pancreas (2μm sections) previously fixed in 4% formaldehyde. Paraffin embedded tissue was as well used for apoptosis analyses, using ApopTag® Red In Situ Detection Kit, following manufacturer’s instructions (cat: S7165-Sigma; Merck, Darmstadt, Germany), and acinar cell proliferation by immunohistochemistry for the nuclear localization of Ki67 (rabbit polyclonal, cat: IHC-00375; Bethyl) (35). For the immunofluorescence experiments, we prepared frozen sections (2μm) of Tissue-Tek® O.C.T. compound-embedded pancreas, which guarantees fast freezing for optimal quality sectioning in a cryostat. The following primary antibodies were chosen: anti-FK2 (cat: BML-PW0150; Enzo life Sciences, Farmingdale, NY), anti-ubiquitin (cat: 43124, Cell Signaling Technology, Danvers, MA, USA), anti-CD68 (cat: ABIN181836, antibody online; Aachen, Germany), anti-Mrc1/CD206 (cat: OASA05048; Aviva Systems Biology, San Diego, CA), anti-Ly6g (cat: 25377, Abcam, Cambridge, MA). The secondary IgG antibodies (fluorophores Cy3 and Alexa488) were purchased from Jackson Immunoresearch (Pennsylvania, USA). The primary antibody dilution variated of 1:200 up to 1:400 with overnight incubation at 4°C and 1:400 for the secondary staining with 2h incubation at room temperature. DAPI (4′,6-diamidino-2-phenylindole; 1:1000) was used as nuclei staining and fluorescent mounting medium (DAKO) to cover the slides before imaging acquisition. Quantification of the CD68 and Ly6G immune staining was estimated manually from 5 random pictures in a magnification of 200-fold from each animal. Total number of cells, through WEKA Segmentation in Image J software, was used to obtain the percentage of positive cells. The same design was applied to calculate apoptosis ratio.

### Isolation of acini

Pancreas were carefully removed from wild-type C57BL/6J, β5i/LMP7^+/+^ (littermate) and β5i/LMP7^−/−^ mice and digested with 1mg collagenase (Collagenase from Clostridium histolyticum; EC.3.4.24.3; SERVA Electrophoresis, Heidelberg, Germany), conditioned in Dulbecco’s modified Eagle medium (DMEM; Thermo Fisher Scientific) with 2% bovine serum albumin and 10mM HEPES, under sterile conditions, as initially described (36). Cell suspension was resuspended in fresh medium without collagenase and after 30 min of resting in water bath at 37 °C, three different approaches were addressed: 1) protease activation was examined in living cells upon an enzyme kinetic reaction up to 60min with 1μM Cholecystokinin (CCK, fragment 26-33; C-2175 Sigma) stimulation whereas necrosis was measured by propidium iodide exclusion (37); 2) acini were directly platted in 12 well plates and incubated with 1μM CCK for 6h or 8h and 24h in a humidified atmosphere of 5%CO_2_ at 37°C, for subsequent total RNA or protein extraction; 3) or challenged with the same dose of CCK for 30min prior addition to the macrophage culture, as described below.

### Co-culture of acini and macrophages

Femur of wild-type C57BL/6J, β5i/LMP7^+/+^ (littermate) and β5i/LMP7^−/−^ mice were collected under sterile conditions. Bone marrow stem cells were isolated by flushing the bone with sterile phosphate-buffered saline (PBS), plated and differentiated into macrophages (BMDM), as previously established (38). At the day of the experiment, BMDM culture received fresh medium, before being exposed to isolated acinar cells for 6 or 9 hours. Supernatant was collected for cytokine measurements and the cells were washed with PBS to withdraw non-adherent cells. Subsequently, cells were harvested with 5mM EDTA in PBS on ice and resulting pellet stored at −80°C for posterior total RNA extraction.

### Transcript expression by quantitative Real-time PCR

All reagents were purchased from Thermo Fisher Scientific, unless otherwise specified. Pancreas, acini and macrophages were collected and total RNA was extracted with Trizol reagent (cat: 15596026), as instructed by the manufacturer using RNAse-free labware. The RNA concentration was measured spectrophotometrically at 260 nm. Next, oligo-dT and random hexamer primer (RH; cat: SO181) were used to generate cDNA from 1-2μg RNA using M-MLV reverse transcriptase (cat: 28025013). Reactions were performed in an Applied Biosystems QuantStudio 7 Flex Real Time System using SYBR Green PCR Master Mix (cat: 4334973), according to the manufacturer’s recommendations. GAPDH or 5S ribosomal RNA expression was used as internal control, detailed primer sequences are shown in the table 1.

**Table 1:**
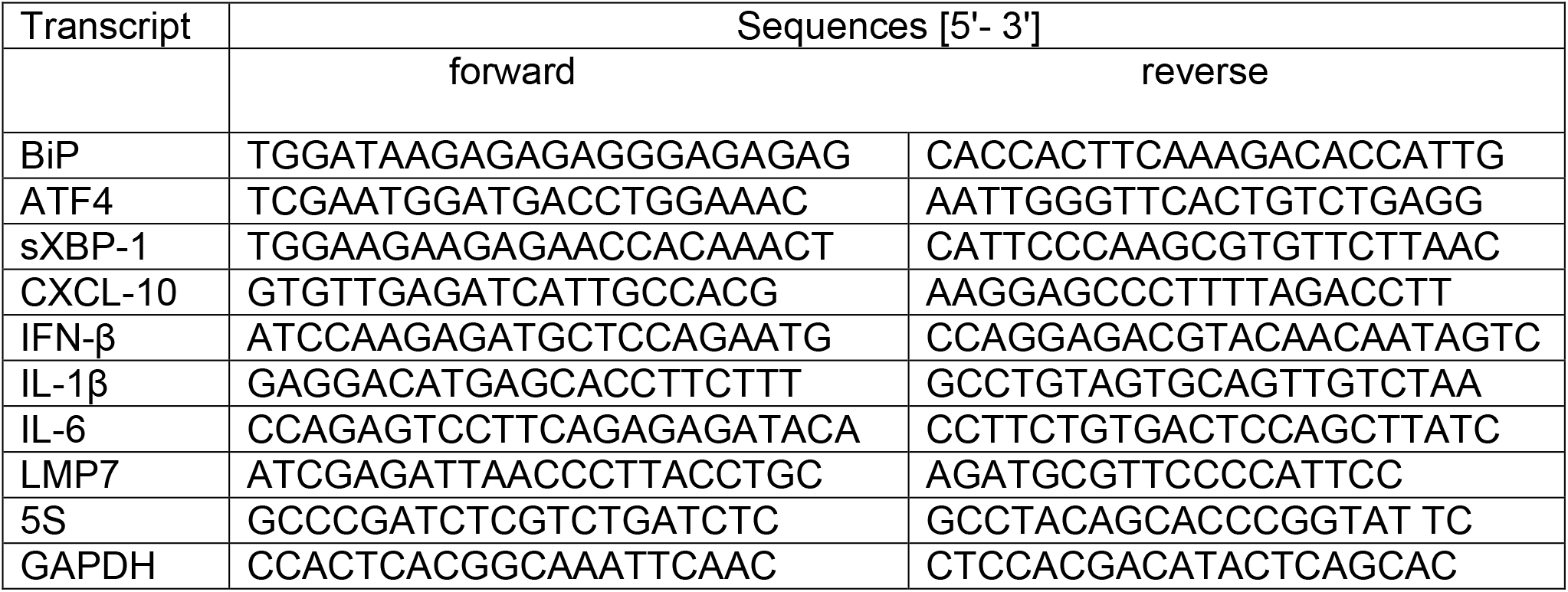
Murine primer sequences for the transcripts assessed by quantitative real-time PCR.

### Measurement of protein levels by western blotting

Protein extracts from pancreas homogenates and acini normalized for protein concentration (10 μg protein) were subjected to SDS–PAGE and then transferred onto PVDF (Polyvinylidene fluoride) or nitrocellulose membranes (Thermo Fisher Scientific) for immunoblotting. Specifically, insoluble ubiquitin-protein conjugates were detected after running 8% gels and PVDF membrane whereas 12% gels and nitrocellulose membrane were used for β5i/LMP7 and CHOP and 15% for LC3-II analyses. Blocking was done with 1% Roti-Block (cat: A151.1; Carl Roth, Karlsruhe, Germany) before primary antibody overnight incubation at 4°C at the following concentrations: anti-LMP7 (1:100000), anti-β5 (1:2000; cat: 3330, Abcam), anti-ubiquitin (1:5000; FK2, BML-PW0150, Enzo Life Sciences), anti-CHOP (1:2000; 2895, Cell Signaling) and anti-LC3 (1:2000; 2775, Cell Signaling). All membranes were incubated with secondary horseradish peroxidase conjugated antibody (1:10000; GE Healthcare, Chicago, Illinois, USA) for 1-2 h at room temperature. Afterwards, the membranes were developed by chemiluminescence with SuperSignal™ West Femto-ECL-substrate (cat: 34095; Thermo Fisher Scientific) and imaging capture was acquired by Fusion equipment. Band intensities were quantified densitometrically using the Image J software (https://imagej.nih.gov/ij/) and normalized by loading control (amylase in case of acini and Ponceau staining for pancreas extracts).

### Statistical analyses

Data are presented as means ± SEM and the number of independent experiments is indicated. Statistical analyses, significant when *P* < 0.05, were performed with GraphPad software (GraphPad, San Diego, CA) using one-way analyses of variance (ANOVA) followed by Turkey’s multiple comparison test or unpaired Student’s *t*-test, as appropriate.

For the Suppl. Fig.1, SigmaPlot 11.0 (Systat Software, Erkrath, Germany) was used for the analyses.

## Abbreviations used in this paper

ATF4: Activating transcription factor 4
BIP: binding immunoglobulin protein
BMDM: bone marrow derived macrophages
CCK: cholecystokinin
CD68 and CD206: cluster of differentiation 68 and 206
CHOP: C/EBP homologous protein
CXCL-10: C-X-C motif chemokine ligand 10
ER: endoplasmic reticulum stress
IFN: interferon
IL: interleukin
LC3: Microtubule-associated proteins 1A/1B light chain 3B
LMP2 and LMP7: Large multifunctional peptidase 2 and 7
Ly6G: Lymphocyte antigen 6 complex locus G6D
MECL-1: multicatalytic endopeptidase complex-like-1
MPO: myeloperoxidase
UPR: unfolded protein response
UPS: ubiquitin protein system
XBP1: X-box binding protein 1

## Acknowledgments

We would like to thank Davide Figini for helping us with imaging analyses and Kathrin Gladrow for excellent technical assistance.

## Figure Legends

**Figure 1: Expression and regulation of LMP7 subunit in isolated acini and in experimental acute pancreatitis**. A) Isolated acini were treated with 1μM of CCK for 8h and 24h. The levels of LMP7 and β5 subunits were determined by western blotting. Amylase was used as an internal control and a ratio of LMP7 was calculated. B) Transcript levels of LMP7 were evaluated by real-time PCR in the pancreas after caerulein-induced pancreatitis. 5S was used as an internal control (n=5-8). C) Western blotting analyses of LMP7 subunit in the pancreas during pancreatitis (n=9-10); Data from independent experiments are expressed as means ± SEM. *p<0.05, **p<0.01, ***p<0.001.

**Figure 2: Higher pancreatic damage in the absence of LMP7**. A) Serum amylase, lipase and D) lactate dehydrogenase activities were measured using specific chromogenic substrates. All enzymes are significantly more active after 8h of pancreatitis in LMP7^−/−^ compared to LMP7^+/+^ group. 0h (n=10-13); 8h and 24h (n=20-25). B) Trypsin and chymotrypsin activities were quantified in the pancreas using corresponding fluorescent substrates. Trypsin and chymotrypsin activities are stronger regulated in the absence of LMP7 at 24h group. 0h (n=3-4), 8h (n=8-13) and 24h (n=5-9). C) Apoptotic bodies were detected by tunel DNA fragmentation assay in paraffin-embedded pancreas and DAPI was used as nuclei staining (n=4-6). Apoptosis ratio was computed based on positive versus total number of cells, normalized by the corresponding LMP7^+/+^ 8h group. D) Histology of pancreatic damage was visualized by hematoxylin & eosin staining of pancreas. The immune infiltrating cells correlates with the extension of necrotic areas and subsequent decrease of unaffected healthy exocrine pancreas. E) Histoscore of disease severity, including necrosis and infiltration. No significant differences were observed after 8h of pancreatitis, nonetheless the pancreas from LMP7^−/−^ mice presented with higher cell damage compared to the littermate group at 24h. Image representative of five independent experiments. Scale bars, 50μm. Size of experimental groups: 0h (n=9-10); 8h and 24h (n=14-17). Data obtained from five independent experiments are expressed as means ± SEM. *p<0.05, **p<0.01, ***p<0.001.

**Figure 3: Increased inflammation in the absence of LMP7**. A) and B) Immunofluorescence in frozen pancreas sections was performed using macrophages (CD68; A) and neutrophil (LY6G; B) markers antibodies. Quantification of positive cells for each leukocyte population was calculated based on the total number of cells, visualized by DAPI nuclei staining (CD206 data not shown). 0h (n=3-4), 8h and 24 h (n=8-9). B) Enhanced neutrophil infiltration in LMP7^−/−^ mice is also reinforced by MPO activity in lung. 0h (n=12-13), 8h (n=19-23) and 24h (n=6-9). C) Transcripts levels of IL-1β, CXCL-10 and IFN-β genes are significantly stronger regulated in the absence of LMP7, which were determined by real-time PCR using specific primers with 5S as housekeeping. LMP7^+/+^ groups-0h (n=4-5), 8h and 24h (n=4-7); LMP7^−/−^ groups-0h (n=4), 8h (n=10-13) and 24h (n=4-9). D) Interleukin-6 levels were measured in serum. Data were normalized to the values of the 8h LMP7^+/+^ group. 0h (n=10-13); 8h and 24h (n=20-25). Scale bars, 20μm. Data are representative of five independent experiments and expressed as means ± SEM. *p<0.05, **p<0.01.

**Figure 4: Impairment of ubiquitinated protein degradation in the pancreatic acini from LMP7^−/−^ mice**. Immunofluorescence in pancreas sections performed using ubiquitin A) alone or C) combined with either CD68, Ly6G or trypsin antibodies. The co-localisation of ubiquitin was exclusively visualized with trypsin antibody, DAPI was used as nuclei staining. Scale bars, 20μm. Image representative of five independent experiments. B) Higher accumulation of insoluble ubiquitin conjugates was also visualized by western blotting and its ratio to a loading control (ponceau staining) was estimated by densitometry. (n=8-12). Data are representative of independent experiments and expressed as means ± SEM. *p<0.05.

**Figure 5: Stronger induction of ER stress transcripts in acute pancreatitis and isolated acini from LMP7^−/−^ mice.** ER stress transcripts levels of BIP, ATF4 and sXBP-1 were determined by real-time qPCR using specific primers and 5S or GAPDH as housekeeping. A) RNA total from pancreas after pancreatitis; LMP7^+/+^ (n=10-13) and LMP7^−/−^ groups (n=10-21). B) CHOP protein expression was assessed by western blotting in the pancreas (n=9-12). C) Isolated pancreatic acini stimulated with 1μM of CCK for 6h (n=3). D) Co-culture of acini and macrophages cells for 6h at 37°C (n=3-5). Data are illustrative of independent experiments and expressed as means ± SEM. *p<0.05; **p<0.01; ***p<0.001.

**Figure S1: Intracellular protease activation and cell death rate in isolated acini.** Trypsin and CTSB activation as well as cellular necrosis were not primarily affected by LMP7 depletion. Acini were supramaximal stimulated with CCK and both enzymatic activities assessed with fluorescent substrates in a time-course reaction. Cell death was measured by propidium iodide inclusion. Data are representative of independent experiments and visualized by means ± SEM. (n=8-10).

**Figure S2: Production of pro-inflammatory cytokines *in vitro*.** The corresponding transcripts were assessed by real-time qPCR. 5S and GAPDH were used as housekeeping for acini and macrophages, respectively. A) Isolated acini were incubated with 1μM of CCK for 6h at 37⁰C. B) Co-exposed acini macrophages were incubated in cell culture for 9h at 37⁰C. Secreted levels of IL-6 and IL-1β cytokines were measured in the supernatant by CBA and ELISA, respectively, and IFN-β transcript expression by real-time qPCR. Data were normalized respective to the LMP7^+/+^ 8h group. Data are representative of independent experiments and expressed as means ± SEM (n=3-7). *p<0.05, **p<0.01, ***p<0.001.

**Figure S3: Absence of LMP7 has no influence on autophagy activation**. LC3 isoforms were detected in the pancreas by western blotting analyses. LC3-II ratio to a loading control (ponceau staining) was calculated by densitometry. No differences were observed in the experimental conditions. Data are representative of independent experiments and expressed as means ± SEM. (n=10-12).

**Figure S4: Acinar cell proliferation is unaltered in LMP7^−/−^ mice**. A) Ki67 proliferation marker was assessed in the pancreas by immunohistochemistry using DAB as substrate (brown-black stain). Hematoxylin accounted for nuclei staining and the positive staining from inflammatory cells was excluded from the quantification. Scale bars, 50μm. Image representative of five independent experiments. Data are expressed as means ± SEM. (n=5-7).

**Suppl. Figure 1:**
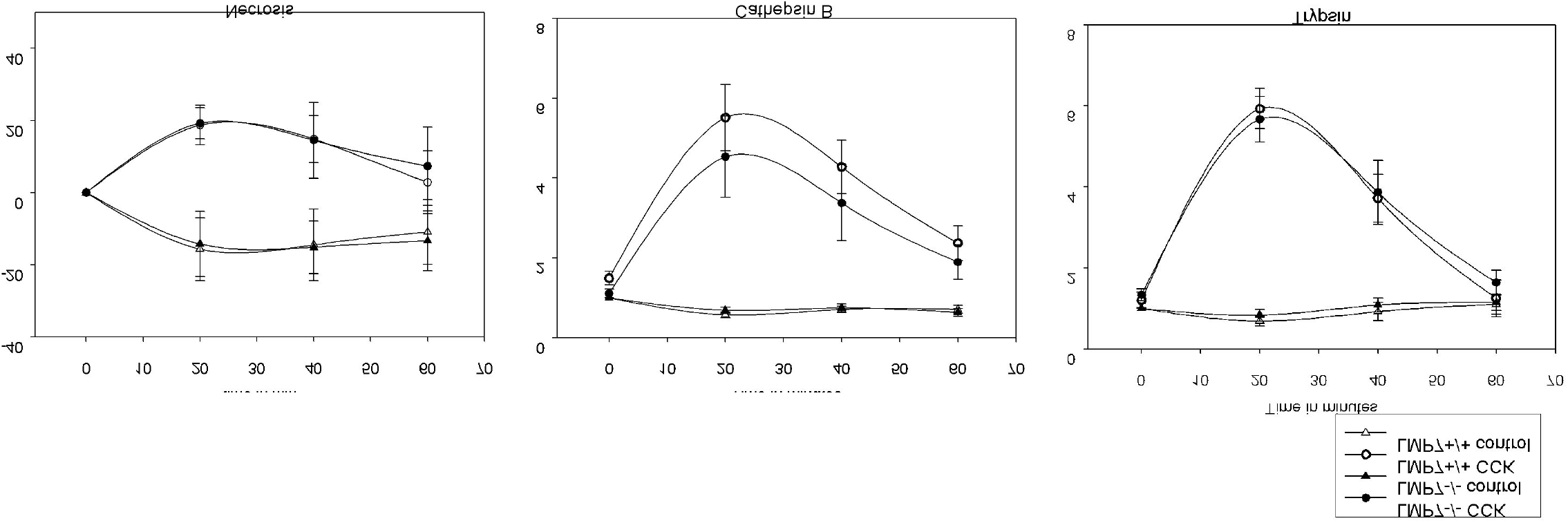
Intracellular protease activation and cell death rate in living acinar cells.

**Suppl. Figure 2:**
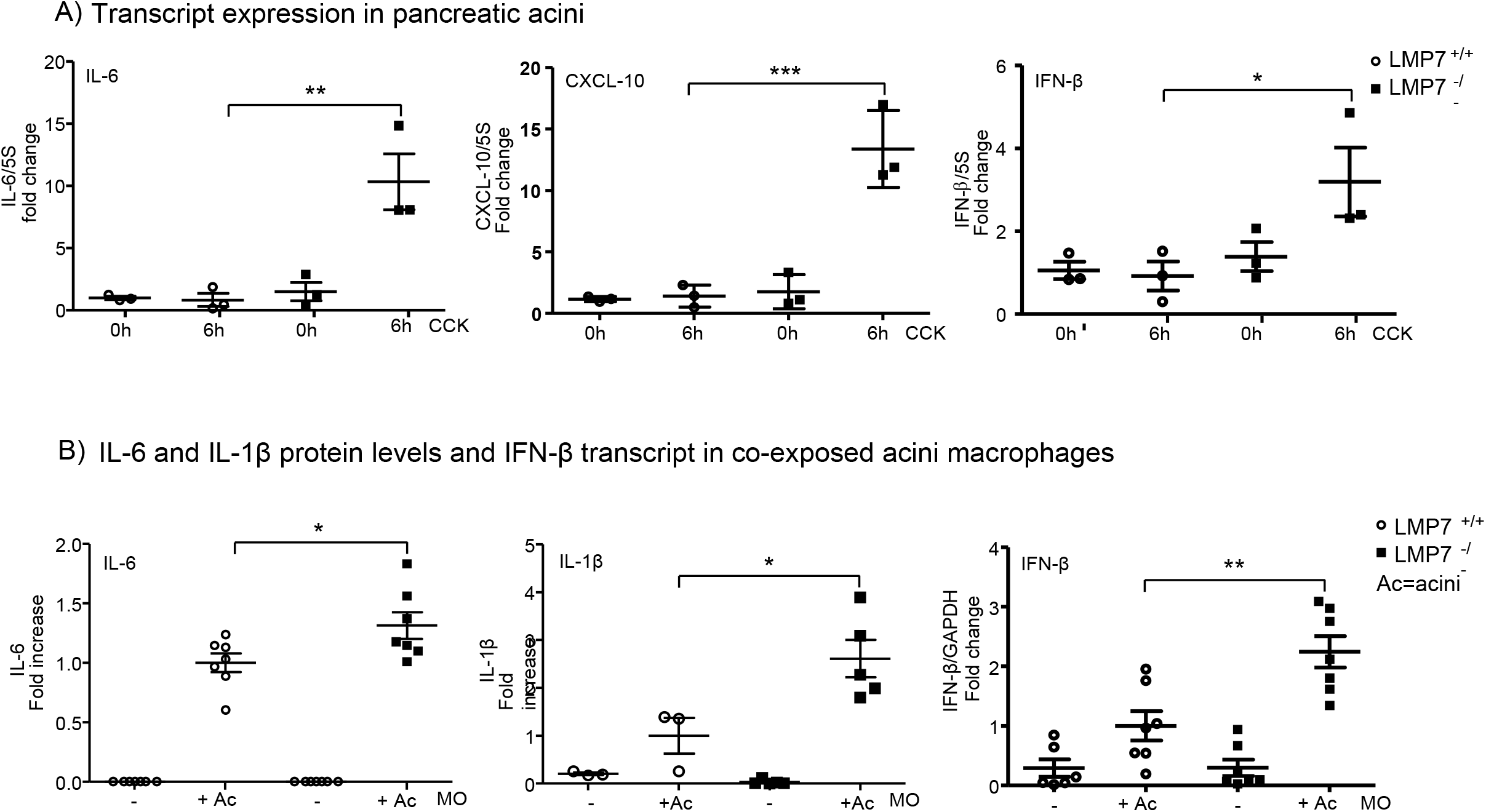
Expression and production of pro-inflammatory cytokines *in vitro*.

**Suppl. Figure 3:**
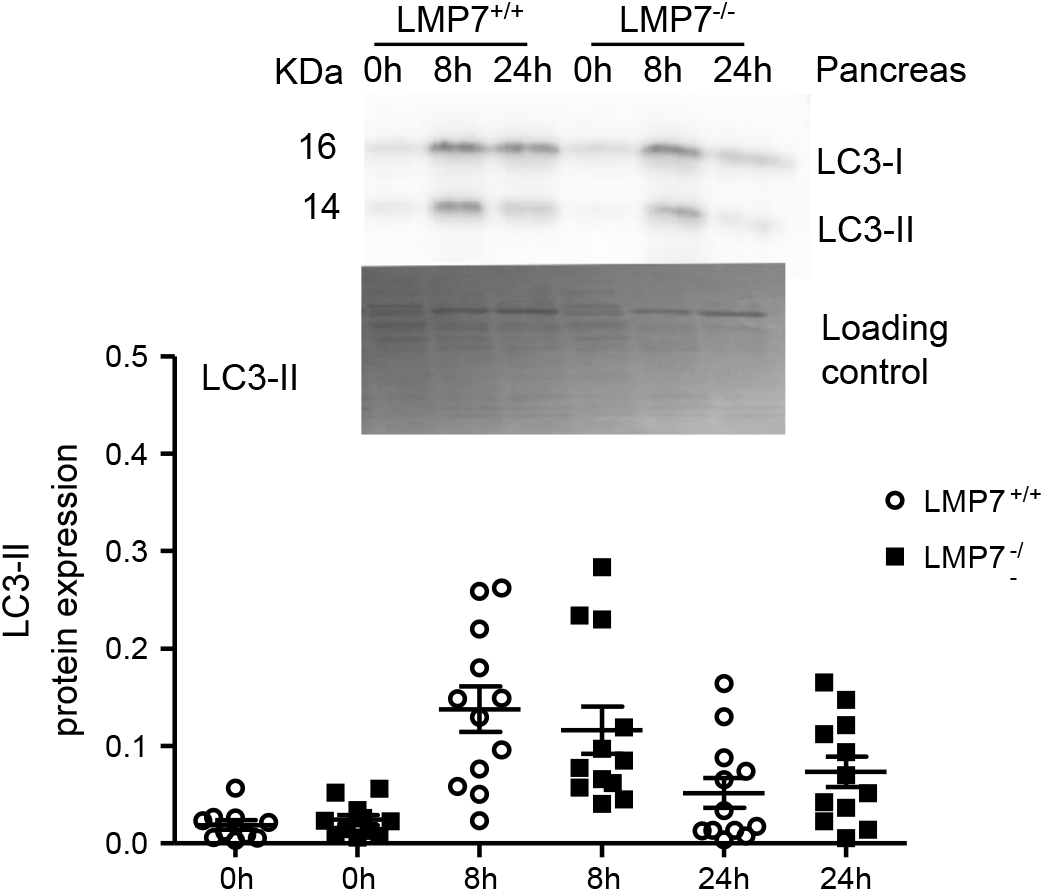
Absence of LMP7 subunit has no influence on autophagy activation.

**Suppl. Figure 4:**
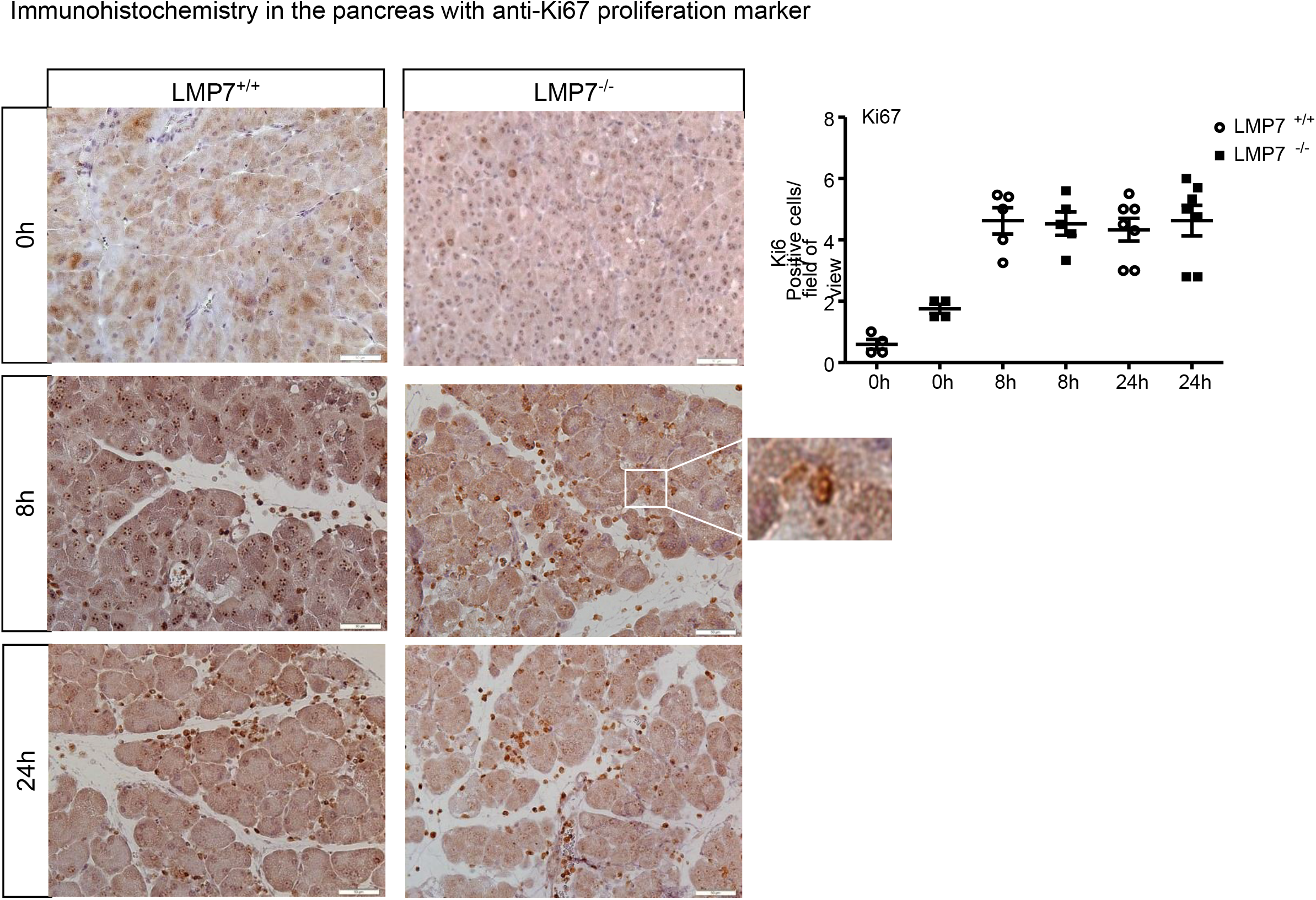
Acinar cell proliferation is unaltered in LMP7^−/−^ mice. Immunohistochemistry in the pancreas with anti-Ki67 proliferation marker

